# Advancing cooperative breeding research with a peer-reviewed and “live” Cooperative-Breeding Database (Co-BreeD)

**DOI:** 10.1101/2024.04.26.591342

**Authors:** Yitzchak Ben Mocha, Maike Woith, Szymon M. Drobniak, Shai Markman, Francesca Frisoni, Vittorio Baglione, Jordan Boersma, Laurence Cousseau, Rita Covas, Guilherme Henrique Braga de Miranda, Cody J. Dey, Claire Doutrelant, Roman Gula, Robert Heinsohn, Sjouke A. Kingma, Jianqiang Li, Kyle-Mark Middleton, Andrew N. Radford, Carla Restrepo, Dustin R. Rubenstein, Carsten Schradin, Jörn Theuerkauf, Miyako H. Warrington, Dean A. Williams, Iain A. Woxvold, Michael Griesser

## Abstract

Research on cooperative breeding (a system with the core characteristic of individuals providing care for the offspring of others) is important for understanding sociality and cooperation. However, large-scale comparative analyses on the drivers and consequences of cooperation frequently use considerably inaccurate datasets (e.g. due to inconsistent definitions and outdated information). To advance comparative research on cooperative breeding, we introduce the <u>Co</u>operative-<u>Bree</u>ding <u>D</u>atabase (Co-BreeD), a growing database of key socio-biological parameters of birds and mammals. First, we describe Co-BreeD’s structure as a (i) sample-based (i.e. multiple samples per species linked to an exact sampling location and period), (ii) peer-reviewed and (iii) updatable resource. Respectively, these curating principles allow for (i) investigating intra- and inter-species variation and linking between fine-scale social and environmental parameters, (ii) accuracy and (iii) continuous correction and expansion with the publication of new data. Second, we present the first Co-BreeD dataset, which estimates the prevalence of breeding events with potential alloparents in 265 samples from 233 populations of 150 species, including 2 human societies (N = 26,366 breeding events). We conclude by demonstrating (i) how Co-BreeD facilitates more accurate comparative research (e.g. increased explanatory power by enabling the study of cooperative breeding as a continuous trait, and statistically accounting for the sampling error probabilities), and (ii) that cooperative breeding in birds and mammals is more prevalent than currently estimated.

## I. Introduction

Cooperative breeding is a system with the core characteristic of individuals providing alloparental care to the offspring of their group members (Brown, 1974; Stacey & Koenig, 1990; Kappeler, 2019; Ben Mocha *et al*., 2023b). Ever since Darwin (1888), this phenomenon has attracted continuous and interdisciplinary research interest (Skutch, 1935; Boland & Cockburn, 2002; Sarano *et al*., 2023). For instance, behavioural ecologists have examined why individuals invest in the offspring of others (Sherman *et al*., 1995; Zahavi & Zahavi, 1997), comparative psychologists have investigated whether alloparental care facilitates the evolution of elaborate social cognition (Burkart & van Schaik, 2010; Tomasello & Gonzales-Cabrera, 2017), and as humans are cooperative breeders, anthropologists have endeavoured to understand cooperative breeding to shed light on the evolution of our species (Hrdy, 2007; Hill & Hurtado, 2009).

Large-scale cross-species comparisons are powerful tools to investigate the causes and consequences of cooperative breeding (e.g. MacLeod & Lukas, 2014; Dey *et al*., 2017; Kingma, 2017; Downing, Griffin & Cornwallis, 2020; Barsbai, Lukas & Pondorfer, 2021; Camerlenghi *et al*., 2022). Using advanced statistical methods that control for phylogenetic relationships, such analyses compare cooperatively versus non-cooperatively breeding species in relation to other parameters of interest [e.g. climatic variables (Jetz & Rubenstein, 2011; Griesser *et al*., 2017) or the prevalence of extra-group offspring (Cornwallis *et al*., 2010; Kingma *et al*., 2010)]. The power of these analyses thus depends fundamentally on the exact definition used to classify species as cooperative breeders and the accuracy of biological parameters ascribed to each species (e.g. annual rainfall in habitat, degree of reproductive skew). Nevertheless, accumulating studies demonstrate that frequently used datasets on cooperative breeding species include numerous errors (Schradin, 2017; Brouwer & Griffith, 2019), significant methodological biases (Sandel *et al*., 2016; Lukas & Clutton-Brock, 2017; Ben Mocha *et al*., 2023a) and vague definitions (Huck, Di Fiore & Fernandez-Duque, 2020; Makuya, Olivier & Schradin, 2022). These shortcomings have resulted in a growing number of scholars questioning the validity of existing datasets (Brouwer & Griffith, 2019; Cockburn, 2020) and calling for more careful data curation (Griesser & Suzuki, 2016; Lukas & Clutton-Brock, 2017; Schradin, 2017).

There are three major sources of inaccuracy in cooperative breeding datasets. First, the use of vague or *ad hoc* definitions of cooperative breeding leads to subjective and inconsistent classification of species as cooperative versus non-cooperative breeders (Lukas & Clutton-Brock, 2017; Ben Mocha *et al*., 2023b). Second, the diversity exhibited by bird and mammal species requires adjusting the way socio-biological parameters are assessed across species. Yet, it is unlikely that any small group of curators could master this manifold diversity to assess parameters in the most biologically meaningful way for each species (section II.1). Third, incorrect information, such as typographic errors (Brouwer & Griffith, 2019), is preserved since paper-associated datasets are rarely corrected and updated.

In addition, most current datasets are limited in scope. First, species are often represented by a single estimate (e.g. from a single population), thereby hindering exploration of within-species variation and yielding the resulting species classification more prone to random or biased sampling errors (e.g. due to small sample size, see also ESM) (Barsbai *et al*., 2021). Second, most comparative datasets focus on one taxon (e.g. birds, nonhuman mammals or humans), thus preventing a more comprehensive study of cooperative breeding. Third, datasets rarely include comparative data from human populations, thereby hampering research about human evolution within the broader theoretical framework of cooperative breeding (Burkart, van Schaik & Griesser, 2017; Ben Mocha, 2020).

Here we introduce the Cooperative Breeding Database (Co-BreeD), a platform designed to standardise and advance the comparative study of cooperative breeding. First, we provide a methodological account of Co-BreeD’s structure and how its curating principles are designed to overcome the above-mentioned limitations. Next, we present a methodological account of the first socio-biological dataset curated to Co-BreeD: the prevalence of breeding events with potential alloparents. Finally, we conclude with a descriptive summary of this dataset and demonstrate how it increases the explanatory power of statistical analyses by enabling the study of cooperative breeding as a continuous trait (e.g. by using the proportion of breeding events with alloparents) and accounting for the probability of sampling error.

## II. Co-BreeD: Cooperative-Breeding Database

Co-BreeD is a growing database covering key biological parameters relevant to cooperative breeding research of bird and mammal species (including humans). It is a multi-dataset database of population-level estimates of biological parameters. The initial dataset curated in Co-BreeD presents data on the prevalence of breeding events with potential alloparents across populations (see section III.3). Additional datasets that are being curated include estimates of within-group female reproductive sharing and allonursing, with plans to curate datasets on other types of alloparental care (e.g. the share of alloparents in feeding and carrying offspring).

### II.1. Important features of Co-BreeD

#### Updatability

Co-BreeD favours a relatively small number of accurate data entries that were peer-reviewed by the original researchers over a larger sample size with potentially less precise estimates. This preference was advocated by multiple behavioural ecologists (e.g. Griesser & Suzuki, 2016; Heldstab, van Schaik & Isler, 2017; Schradin, 2017) and statisticians (Meng, 2018; Bradley *et al*., 2021), with the latter demonstrating that more accurate results were obtained from smaller samples of accurate data than from larger samples of less accurate data. Since the care system of most bird and mammal species is unknown or indirectly inferred (Cockburn, 2006; Griesser & Suzuki, 2016; Lukas & Clutton-Brock, 2017), curating a reliable database of the care systems of all species of birds and mammals is currently impossible. Co-BreeD is therefore designed as a “living” resource to which new data is constantly being added (e.g. following the publication of new natural history data) and corrections to existing records can be made. To this end, Co-BreeD is deposited in an updatable and open-access GitHub repository.

#### Included species

Resources dependent, Co-BreeD will work to include all bird and mammal species with reliable data on alloparental care (i.e. cooperative breeding species) or parental care (i.e. non-cooperative breeding species). However, at the current stage, Co-BreeD focuses on species reported to exhibit alloparental care. This focus makes Co-BreeD a state-of-the-art resource for studying variability among cooperatively breeding species. Yet, to enable comparison between cooperatively and non-cooperatively breeding species, 12% of the species currently included in Co-BreeD are not regarded as cooperative breeders (i.e. to the best of our knowledge, these species do not exhibit a systematic and unequivocal form of alloparental care as allonursing, allofeeding, offspring carrying and incubation by other group members than the offspring’s biological parents. For instance, leopard *Panthera pardus* and black catbird *Melanoptila glabrirostris*). The species currently included in Co-BreeD were selected due to the availability of relevant data and their representation of diverse taxonomic groups, geographic distribution and social systems (Kappeler, 2019).

#### Sample-based structure

Co-BreeD has a sample-based structure in which each record (i.e. a row in the datasheet) represents a specific sample from a specific population of a species. The meta-data of each sample (Table 1) indicates its exact sampling location and period. In addition, many species are represented with multiple populations and/or multiple sampling periods of the same population. This sample-based structure enables two important features: (i) retrieving fine-scale environmental data for each sample; and (ii) investigating within-species spatial variability and within-population temporal variability.

**Table 1.** Columns of the Sample Meta-Data section (SMD)

| **Table 1. Columns of the Sample Meta-Data section (SMD)** |  |
| --- | --- |
| **Column header** | **Explanation** |
| taxonomic_class_SMD | The taxonomic names follow the Handbook of the Birds of the World and BirdLife International Checklist (2022: version 7) for birds and the Mammal Diversity Database (2022: version 1.11) for mammals. |
| taxonomic_order_SMD |  |
| species_english_name_SMD |  |
| species_scientific_name_SMD |  |
| culture_name_SMD | As described in the publication from which the data were extracted (only for humans). |
| species_level_evidence_for_alloparental_care_by_Ben_Mocha_et_al._2023_definition_SMD | Categorical classification of whether, to the best of our knowledge, the species exhibits sufficient evidence for a systematic and unequivocal form of alloparental care in >5% of breeding events in $\geq 1$ population (i.e. suspected yes / yes / no evidence for nursing, feeding, incubation, or carrying of offspring by alloparents, see Ben Mocha *et al.* (2023b) for discussion). This parameter can be used as a suggestive list of Co-Breed species that are qualified as cooperative breeders. This list should be used with caution as it is not based on a comprehensive review of each species. |
| sampling_location_SMD | A verbal description of the name of the sampling location. The description ends with “, the country name”. |
| latitude_SMD | The coordinates of the sampling location. For uniformity, all coordinates were extracted from Google Maps (https://www.google.com/maps/) and are in the World Geodetic System 1984 (WGS84) standard. |
| longitude_SMD |  |
| start_sampling_period_SMD | The years when sampling began and ended. |
| end_sampling_period_SMD |  |
| unsampled_years_SMD | Unsampled years between the start and end sampling period. |
| reference_title_SMD | The title of the publication. |
| DOI_SMD | The DOI of the publication. |
| comments_SMD | Comments regarding any of the Sample Meta-Data (SMD) columns. For example, when the indicated coordinates are only approximations of the sampling location. Comments can be made by Co-Breed curators and/or the authors who confirmed their data. |

In addition, when available, Co-BreeD curates a detailed year-by-year date for samples as an appendix of each dataset on a specific biological parameter.

#### Reviewing and archiving unpublished data

Co-BreeD does not accept unpublished data unless (i) the data owner published the methods used to obtain these data elsewhere, (ii) Co-BreeD curators reviewed the unpublished data, and (iii) the data are published as a Co-BreeD year-by-year appendix. In archiving unpublished datasets from habitats that have been substantially modified or are at risk of human modification, Co-BreeD also helps to preserve knowledge about the behavioural adaptations that evolved for living in these natural environments (Kühl *et al*., 2019).

#### Objective classification tools

Co-BreeD offers two unique classification capabilities. First, it facilitates a more accurate binary classification of species as cooperatively versus non-cooperatively breeding species. By curating biological parameters that are considered defining conditions for cooperative breeding (i.e. a minimum prevalence of alloparents and systematic occurrence of alloparental care), Co-BreeD users can choose their definition of cooperative breeding and filter species that fulfil the combination of biological criteria required by this definition. For instance, it will allow filtering for species with >10% of breeding events with alloparents (Cockburn, 2006) and/or female breeding monopolisation (Federico *et al*., 2020; Clutton-Brock, 2021). This pluralistic, yet objective, tool for binary species classification can also be used to test whether the conclusions of a comparative study depend on the used definition. We advise users who apply a binary species classification to choose a specific cooperative breeding definition and carefully filter those species that fulfil its criteria. Table 1 provides details about the list of Co-BreeD species suggested to be considered as cooperative breeders.

Second, by providing continuous estimates of biological parameters, Co-BreeD enables the study of cooperative breeding as a continuous trait (e.g. using the percentage of breeding events with alloparents in the population as the response variable). For example, instead of a binary classification of the black catbird as a non-cooperative species, and the long-tailed tit (*Aegithalos caudatus*) and greater ani (*Crotophaga major*) as cooperative species, the “degree” of cooperative breeding in these species can be described by the percentage of breeding events with alloparents in a population [0% of broods attended by alloparents in a black catbird population in Mexico (Lapergola, 2012), 54% of broods attended by alloparents in a long-tailed tit population in the UK (Hatchwell & Russell, 1996) and 100% of broods attended by alloparents in a greater ani population in Panama (Riehl & Smart, 2022)]. Studying cooperative breeding as a continuous trait accounts for the ample variation in the occurrence of alloparental care across and within cooperative breeding species (Figure 1) and provides considerably more explanatory power over binary classification (Clutton-Brock, 2021; Olivier *et al*., 2024).

**Figure 1.**
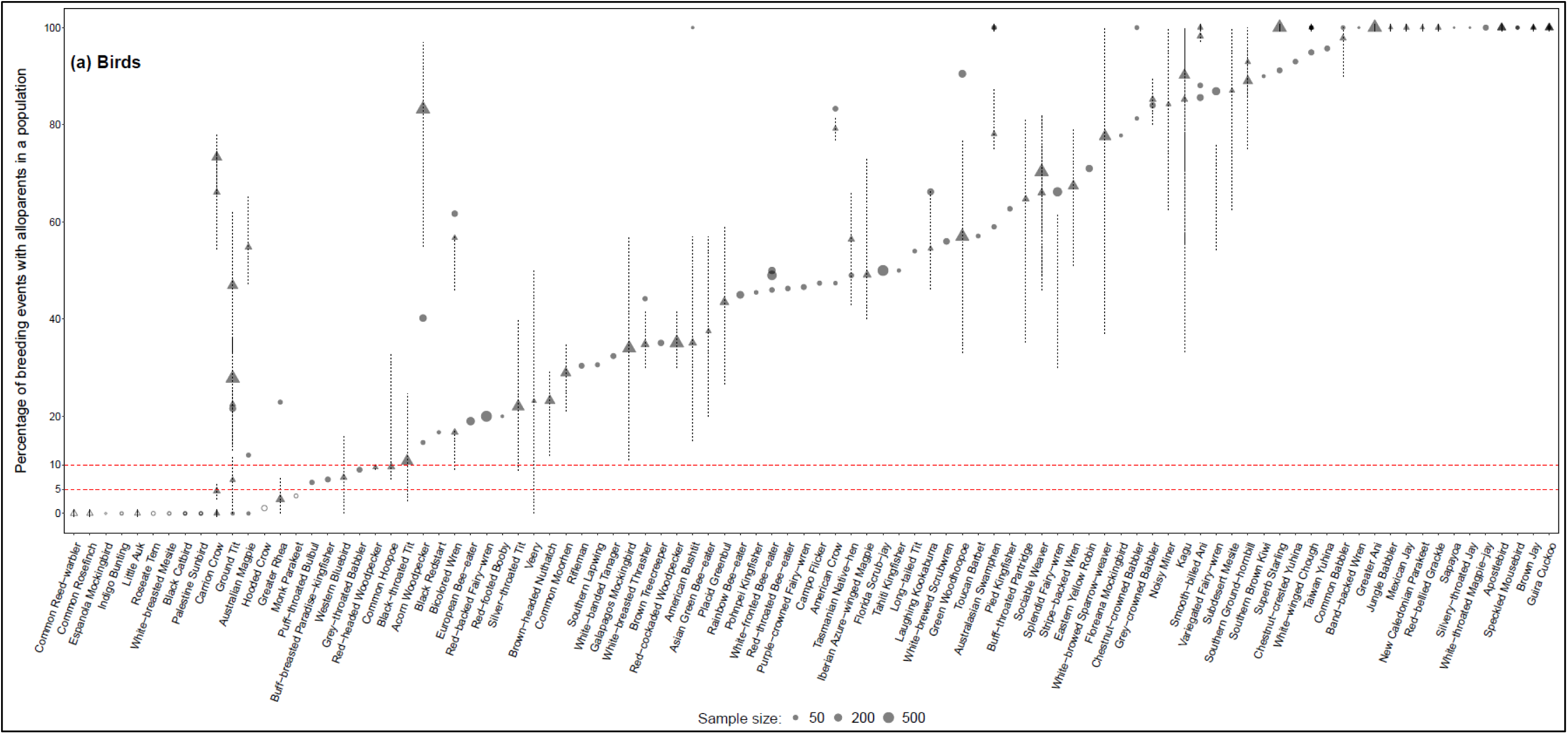
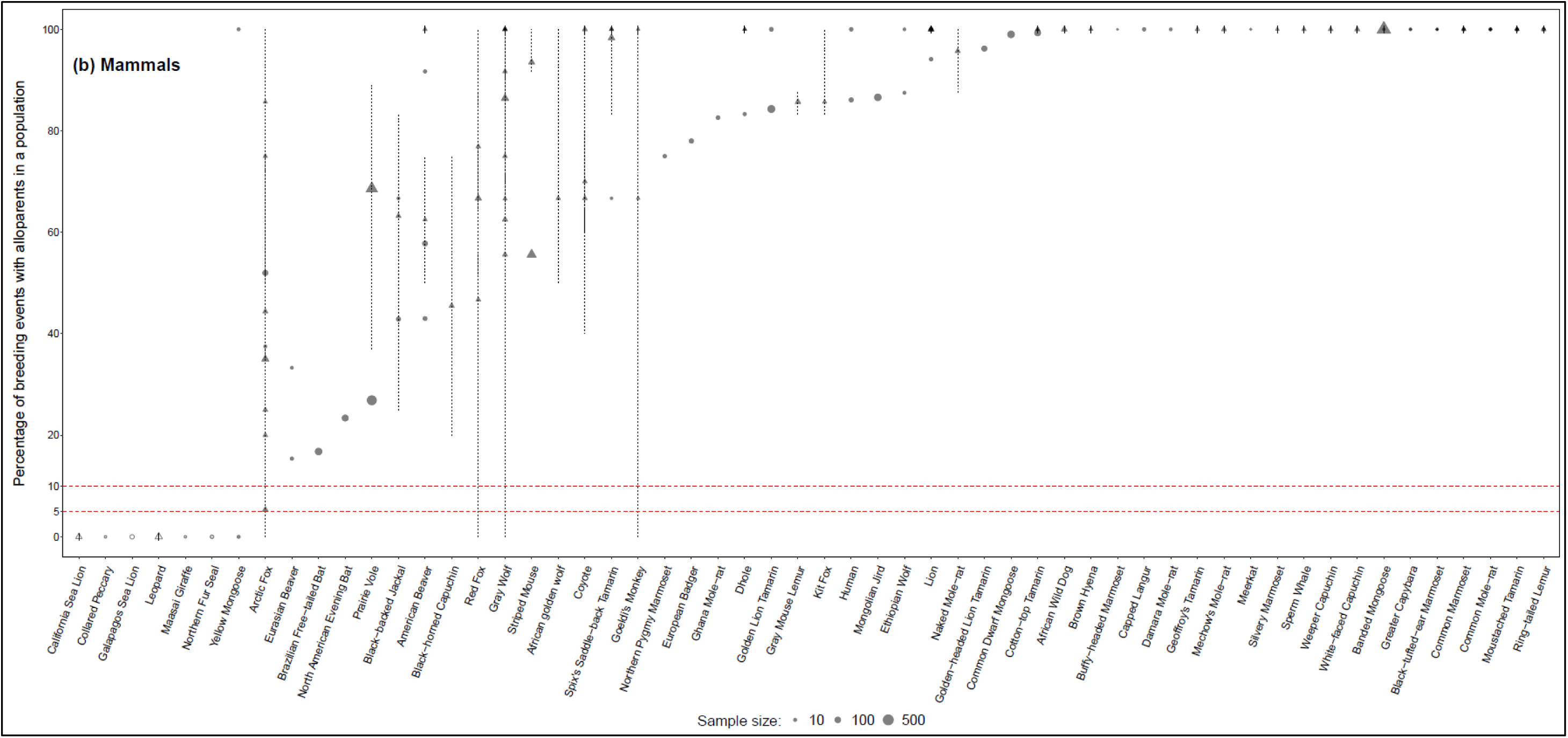
Estimates of the proportion of breeding events with potential alloparents in different samples of (a) birds (N = 147 samples, 132 populations, 94 species) and (b) mammals (N = 118 samples, 101 populations, 56 species) included in the Co-BreeD *PA* dataset. Species are presented in ascending order starting from the population with the lowest prevalence of breeding events with potential alloparents. Circles represent the percentages of breeding events with potential alloparents during a specific period and populations. Triangles within lines represent the percentage and multi-year range (respectively) of breeding events with potential alloparents of the same population. The size of symbols reflects sample size (limited to 500 to display a reasonable size of samples of low size). Darker shapes represent overlapping estimations from multiple samples. Red lines represent the minimum thresholds to consider a species as a cooperative breeder as suggested by Cockburn (2006) / Ben Mocha *et al*. (2023b) (i.e. >10% / 5% of breeding events with alloparents in at least one population of the species, respectively). Empty shapes represent samples from species that, to the best of our knowledge, have no population that exhibits a systematic and unequivocal form of alloparental care (e.g. allonursing, allofeeding, carrying of offspring and incubation, see Table 1 for details). For clarity, in samples with a multi-year range without variation (e.g. 0% across all years), the minimum and maximum ranges were manipulated to be one percentage lower and higher, respectively (e.g. −1% – 1% instead of 0-0% in Leopard). These manipulated ranges are represented by solid lines.

#### Statistically accounting for the sampling error probabilities

For the majority of species, data on alloparental and parental care is based on a small sample size (Cockburn, 2006; Schradin, 2017) and is thus prone to relatively high sampling error rates. Co-BreeD minimises this problem by providing the sample size from which each estimate was calculated, thereby enabling giving each estimate in the analysis a different weight according to its sample size (Figures 1 and 2). See ESM for a demonstration of power analysis using Co-BreeD data.

**Figure 2.**
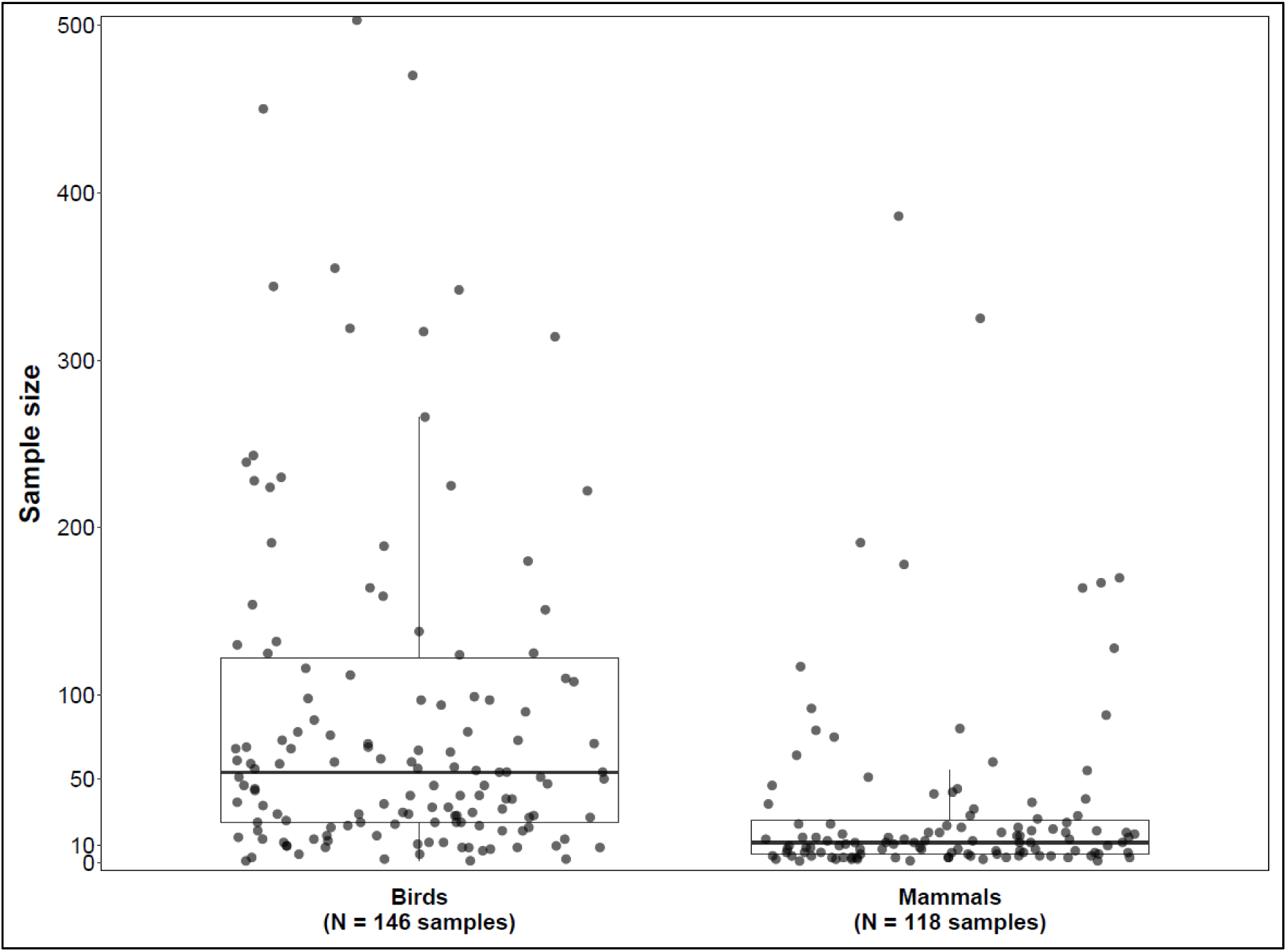
The sample size upon which the estimations of the Co-BreeD *Prevalence of Alloparents* dataset are based. Horizontal bars represent the median, boxes the 25% and 75% quartile and vertical lines indicate the minimum and maximum values. Each jittered circle represents a sample. For clarity, eight and one maximal outliers were excluded from the bird and mammal samples, respectively (range: 503 – 3,234 breeding events).

#### Peer review and adjustment to species-specific characteristics

In-depth familiarity with the social system of a species is important as nuances may require different ways to calculate socio-biological estimates (Lukas & Clutton-Brock, 2017; Brouwer & Griffith, 2019). For example, the basic social unit from which reproductive skew indexes should be calculated may differ according to whether the species lives in discrete groups (e.g. Arabian babblers *Argya squamiceps*, whereby only the members of the discrete social group should be considered when calculating reproductive skew: Lundy, Parker & Zahavi, 1998) or multi-level societies consisting of multiple sub-groups (e.g. bell miners *Manorina melanophrys*, where all members of the higher-level social tier should be considered when calculating reproductive skew: Painter *et al*., 2000; Papageorgiou & Farine, 2021). To ensure that biological estimates are assessed in the most biologically meaningful way for each species, Co-BreeD curators ask the authors of the original publications to confirm the data entry that is based on their publication.

#### Focus on data from natural populations

Co-BreeD estimates natural variation in alloparental care. It thus only includes data from free-living populations, but not captive populations or data from experimental studies in which the experimental manipulation may affect parental care.

#### Quality control

To ensure accuracy and to minimise the risk of transcription errors (e.g. typos), Co-BreeD has a three-step data entry protocol (Figure 3). First, each biological estimate is calculated from the primary literature (reviews are not considered) by a Co-BreeD curator. Second, the data entry is checked by another Co-BreeD curator. In the third step, the author of the publication from which the estimate was calculated is asked to confirm this entry.

**Figure 3.**
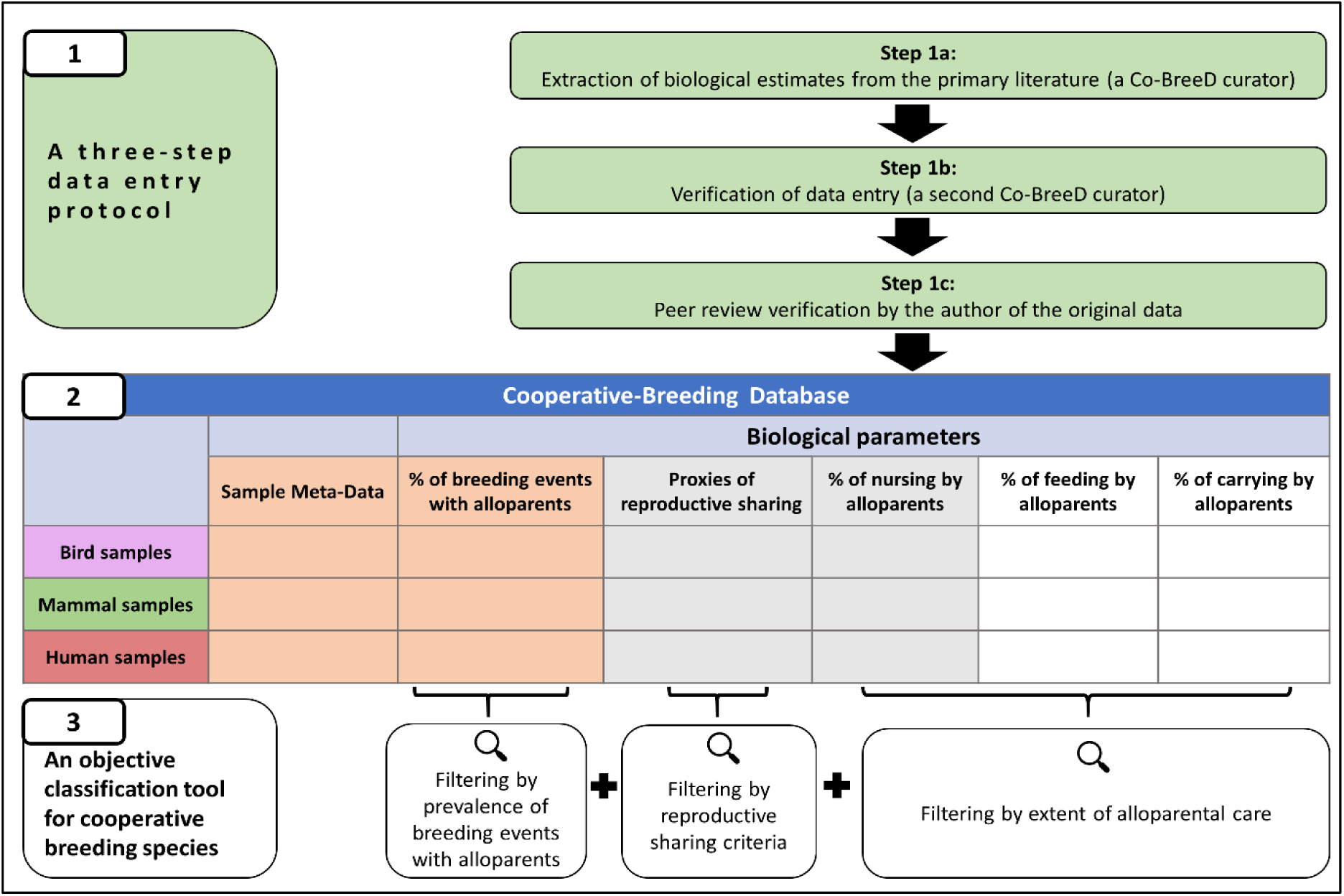
The structure of the Cooperative-Breeding Database. **(1)** The three-step process of data extraction and verification. **(2)** Compiling a sample-based database of key biological parameters. **(3)** Co-BreeD enables filtering species according to the conditions set by different cooperative breeding definitions. The orange-coloured biological parameters are the Co-BreeD sections published in this paper, the grey-coloured parameters are sections that are under preparation and the white-coloured parameters are planned sections.

#### Reproducibility

The reproducibility of the curated data is ensured in two ways. First, we provide a reference including the exact page numbers where the relevant data for the calculated biological estimates are indicated. Second, the methods used to curate each biological parameter will be described in a methodological paper, for example, section III of this paper describes the methods used to curate the dataset on the prevalence of potential alloparents.

#### Fairness

Co-BreeD acknowledges the importance of authors verifying their data and contributing unpublished datasets. Authors who contribute to Co-BreeD are invited to co-author the methodological account of the biological parameter they contributed to or its future updates.

### II.2 The structure of Co-BreeD

Co-BreeD is provided as an Excel file with multiple sheets and separate text-only CSV files generated from each tab for maximum compatibility. The main sheet is named “Biological Estimates” and it presents the different biological parameters for each sample (e.g. the prevalence of breeding events with alloparents and the prevalence of allonursing observed in each sample). Additional sheets in the data file are (i) an “Index” sheet with explanations for each column head, and (ii) appendix sheets with year-by-year data, each for a different biological parameter curated to Co-BreeD.

Having a sample-based structure, each data row in the Biological Estimates sheet depicts a sample from a specific population and study period (in years). Each data row has multiple sections. The first section is a cluster of 15 columns presenting the Sample Meta-Data of each sample. Each of the remaining sections includes a cluster of columns presenting a dataset of a specific biological parameter retrieved from this sample (currently there is one more section presenting data about the prevalence of breeding events with potential alloparents, see section III). To facilitate data processing, each header has a suffix according to the section it belongs to. The Sample Meta-Data columns have the suffix “SMD” and are explained in Table 1.

## III. Co-BreeD *PA* dataset: Prevalence of breeding events with potential alloparents

Even if a species exhibits alloparental care, this behaviour often does not occur in all breeding events within a specific population or in all populations (Figure 1). This variation introduces two problems for comparative research that uses binary species classification as cooperative vs. non-cooperative breeders. First, binary species classifications ignore the ample inter- and intra-species variation in the prevalence of breeding events with alloparents (Figure 1). The binary species classifications, thus, lose considerable explanatory power (Clutton-Brock, 2021; Olivier *et al*., 2024). Second, in well-studied species, even a rare behaviour is likely to be observed (Price, Millington & Grant, 1983; Nichols & Arbuckle, 2022) or mistaken interpretation to be reported (e.g. potential alloparental care in Eurasian coot *Fulica atra*: Carr, 1993; *mute swan Cygnus olor*: Włodarczyk & Kołaciński, 2001). However, if a single case of alloparental care is sufficient to classify a species as a cooperative breeder, species will be systematically false-positively classified as cooperative breeders with the increase in research effort. The binary species classification is, therefore, a one-way process in which species previously classified as non-cooperative breeders can only be false-positively re-classified as cooperative breeders, but not the other way around (Griesser & Suzuki, 2016; Ben Mocha *et al*., 2023b). To facilitate a solution to this problem and enable a greater explanatory power than binary classification, the first Co-BreeD dataset provides quantitative estimates of the prevalence of breeding events with potential alloparents as a continuous measure of cooperative breeding.

### III.1. Methods

#### Data collection

We used Ben Mocha and colleagues’ (2023b) dataset as a starting point to curate the *Prevalence of Alloparents* (*PA* hereafter) dataset. Data on additional species are constantly being added by reviewing the literature on species classified as exhibiting alloparental care by other datasets on birds (Cockburn, 2006) and mammals (Isler & van Schaik, 2012; Heldstab *et al*., 2017). For each species, we collect data on as many populations as possible by searching Google Scholar, ResearchGate and Web of Science.

#### Breeding events with potential alloparents

To estimate the prevalence of breeding events with potential alloparents in a sample, we calculated the percentage of *“breeding events with potential alloparents”* out of the total number of breeding events sampled. Below, we discuss what we considered “breeding events”, “potential alloparents” and how non-cooperative breeding species were assessed.

#### Breeding events

Data permitting, we considered each brood/litter as a breeding event. This means that a social group with multiple breeding females was counted according to the number of broods/litters produced by all the females in the group. Focusing on the broods/litters level results in fine resolution of the prevalence of breeding events with potential alloparents, especially in species where only some broods/litters within the social group receive alloparental care (e.g. Baldovino & Di Bitetti, 2008; Pike *et al*., 2019).

We present examples of how breeding events were considered in three common social systems. First, in family-based groups, often there is a single breeding female (e.g. southern pied babblers *Turdoides bicolor*; Figure 4a), and in the absence of detailed data about shared maternity, we considered each brood as one breeding event. Second, in species living in groups with multiple breeding females but without discrete breeding units, we considered the brood/litter of each mother as a breeding event (Figure 4b). For instance, in the multi-male-multi-female groups of white-faced capuchins *Cebus capucinus* (Sargeant *et al*., 2015), we calculated the number of infants receiving alloparental care out of the total number of infants observed (in this monotocous species, each infant represents a breeding event). To allow reproducibility of how estimates were calculated for each sample, we state what we considered as breeding events with potential alloparents for this sample (Table 2). Third, in plural breeding species and species living in multi-level societies, multiple breeding pairs or “breeding units” are clustered together in a larger social structure (Figure 4c). In these species, we considered each brood/litter from each breeding pair or unit in the group as a breeding event. Examples include superb starlings *Lamprotornis superbus* (plural breeders), where the greater social group has multiple nests simultaneously (Rubenstein, 2007) and multi-level, small-scale human communities that consist of discrete families (Grueter, Chapais & Zinner, 2012). In such cases, we calculated the percentage of nests/families receiving alloparental care rather than the percentage of social groups/villages with alloparents, respectively.

**Figure 4.**
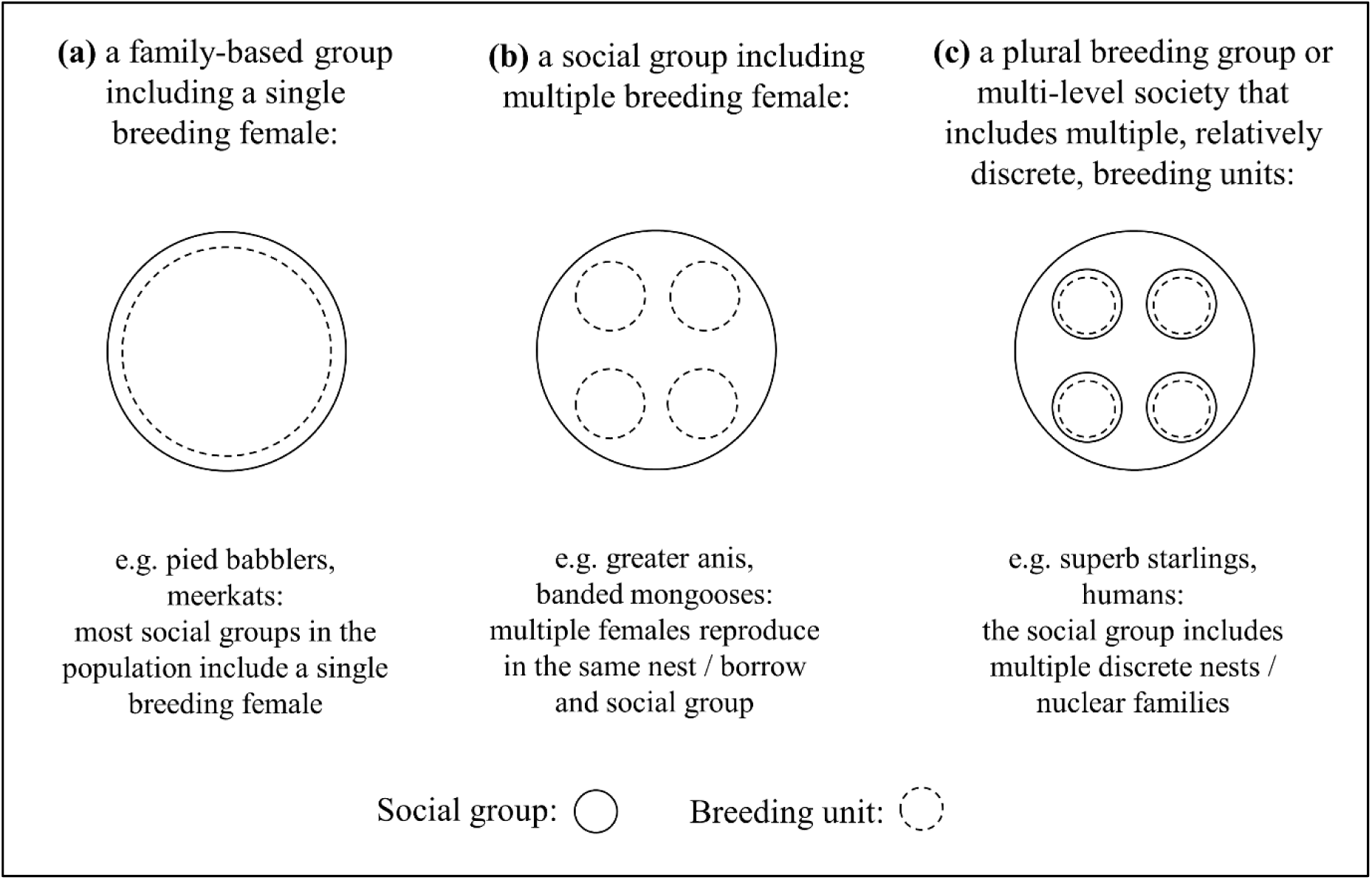
Three common examples of social and breeding structures. The examples present species in which these social and breeding structures are most common, although different combinations often co-occur within a population. Dashed circles indicate the breeding units used to calculate the proportion of breeding events with potential alloparental care.

**Table 2.** Columns of the *PA* dataset (PA)

| **Table 2. Columns of the PA dataset (PA)** |  |
| --- | --- |
| **Column header** | **Explanation** |
| location_of_relevant_data_in_reference_PA | The page number, table number, figure number or supplementary materials with the relevant information. |
| are_year_by_year_data_available_PA | yes / no |
| location_of_year_by_year_data_PA | The page number, table number, figure number, supplementary materials or an unpublished dataset that the authors provided to Co-Breed. Year-by-year data calculated are presented in the PA appendix. |
| data_entry_PA | These columns present the names of the Co-Breed curator who calculated the estimate from the literature, the Co-Breed curator who verified this data entry and the author of the original data who verified the data entry, respectively. The Index sheet provides the full names of the Co-Breed curators. Since Co-Breed is a live resource, the verification status of some entries may be “in process”. Samples for which the authors could not be contacted are indicated as “author could not be contacted”. |
| data_verification_PA |  |
| author_verification_PA |  |
| what_considered_a_breeding_event_with_potential_alloparents_PA | See details in section III.1 |
| percentage_of_breeding_events_with_potential_alloparents_out_of_the_total_number_of_breeding_events_sampled_PA | The percentage of breeding events with potential alloparents out of the total number of breeding events sampled during the entire sampling period.<br>$(\text{number of breeding events with potential alloparents\_PA}) / (\text{sample size\_PA}) = (\text{percentage of breeding events with potential alloparents out of the total number of breeding events sampled\_PA}).$ |
| number_of_breeding_events_with_potential_alloparents_PA | The number of breeding events with potential alloparents sampled during the entire sampling period. |
| sample_size_PA | The total number of breeding events sampled during the entire sampling period. |
| type_of_sample_size_unique_breeding_units_OR_repeated_measurements_PA | Whether the sample size represents unique breeding units (e.g. social groups) or repeated measurements of the same breeding units across multiple breeding events. |
| min_range_<br>the_minimum_yearly_percentage_across<br>_the_sampling_period_PA | These columns give the lowest and highest yearly percentage of breeding events with potential alloparents during the sampling period. Note, that the yearly range could only be calculated for samples longer than one year. For species in which parental care lasts more than a year (e.g. sperm whales *Physeter macrocephalus*: Sarano *et al.*, 2023), we considered each infant-year as a breeding event when calculating the range (contrary to the overall percentage of breeding events with potential alloparents, for which each brood/litter was only considered once). |
| max_range_<br>_the_maximum_yearly_percentage_across<br>_the_sampling_period_PA |  |
| comments_PA | Provides comments regarding the PA dataset, including any caution that needs to be taken when considering this sample. For example, the reported percentage may be underestimated when only nests in which alloparents were observed feeding the nestlings were considered to have alloparents, but limited observations on nestling provisioning were made (e.g. Hughes *et al.*, 1996; Li *et al.*, 2012). |

#### Potential alloparents

Following the distinction made between the parental care system and mating system (Kappeler & van Schaik, 2002; Kappeler, 2019), we consider individuals as alloparents if they provide care to the offspring of other group members, regardless of whether the alloparent breeds at the same time or not. Consequently, we use the term alloparents instead of helpers to avoid implying that individuals who take care of others’ offspring are necessarily non-breeding helpers.

To exclude cases of caring for extra-group offspring, as in brood parasitism, we only consider cases where alloparents care for the offspring of their group members (see discussions in Griesser & Suzuki, 2016; Ben Mocha *et al*., 2023b).

We only considered breeding events as having potential alloparents if typical alloparents of the focal species were present in the social group. For instance, in species where females do not provide alloparental care, only multi-male groups were considered as having potential alloparents.

We present data on breeding events with “potential” alloparents to acknowledge that alloparental care is sometimes assumed according to specific criteria and not directly observed. For example, based on observations that non-parent group members often provide alloparental care in a given population, some studies considered broods in groups with more than two adults as breeding events with potential alloparents (Baglione, Marcos & Canestrari, 2002; Roldán *et al*., 2013). In addition, when data on the number of broods/litters was not available, we used the number of breeding females in a group-year as a proxy of breeding events for this group-year (because in birds and mammals all broods/litters are produced by females, including those with shared paternity). This approach may mix group-living and cooperative breeding, especially in species living in groups but that rarely engage in alloparental care, such as ring-tailed lemurs (*Lemur catta:* Gould, 1992) and buff-throated partridge (*Tetraophasis szechenyii*: Xu *et al*., 2011). Estimates that are based on group size alone should therefore be treated with caution even if the species is known to engage in alloparental care. These group-level estimates will be replaced when more fine-tuned samples of the species become available and can be identified using a dedicated column in the dataset (Table 2).

#### Assessment of non-cooperatively breeding species

To allow comparison between cooperative and non-cooperative breeding species (i.e. species not known to exhibit systematic alloparental care, for example leopards), we needed a conceptual framework that would have allowed detecting the presence of alloparental care if it had occurred in these species. To this end, we defined breeding events with potential alloparents as those that would hypothetically be considered as having alloparents if observed. For instance, >1 mother associated with the cubs in leopards (Balme *et al*., 2017) or >2 birds feeding the nestlings in Palestine sunbirds *Cinnyris osea* (Markman, Pinshow & Wright, 2002) would have been considered as litters/broods with alloparents if observed. We then reviewed studies on at least one form of parental care under the assumption that an unusual behaviour as non-parents caring for the offspring would be reported if observed. If alloparenting was not reported, we assumed that none of the observed breeding events had alloparents (only studies on individually recognised animals were considered).

Since the *PA* dataset estimates the prevalence of breeding events with alloparents in a sample, we did not consider studies reporting breeding events with alloparents without the baseline prevalence of breeding events without alloparents (e.g. case reports of unusual behaviour: Carr, 1993; Nichols & Arbuckle, 2022).

### III.2. The structure of the Co-BreeD *PA* dataset

The Co-BreeD *Prevalence of Alloparents* dataset consists of 14 columns. The title of each column ends with the suffix “PA” and is explained in Table 2.

### III.3. Results and discussion

#### Sample characteristics

The *PA* dataset currently includes 265 samples documenting over 26,366 breeding events of 150 bird and mammal species, including two human populations (Table 3).

**Table 3.** Sample size of Co-BreeD *Prevalence of Alloparents* dataset (as of April 2024).

| Table 3. Sample size of Co-BreeD *Prevalence of Allopurents* dataset (as of April 2024). |  |  |  |  |
| --- | --- | --- | --- | --- |
|  | Number of species | Number of populations | Number of Samples | Samples confirmed by an author of the original study |
| **Birds** | 94 | 132 | 147 | 35% |
| **Non-human mammals** | 55 | 99 | 116 | 10% |
| **Humans** | 1 | 2 | 2 | 50% |
| **Total** | 150 | 233 | 265 | 24% |

Geographically, Co-BreeD *PA* samples are distributed across six continents (Figure 5), with the exception of the absence of bird samples from Antarctica and mammal samples from Australia and Antarctica.

**Figure 5.**
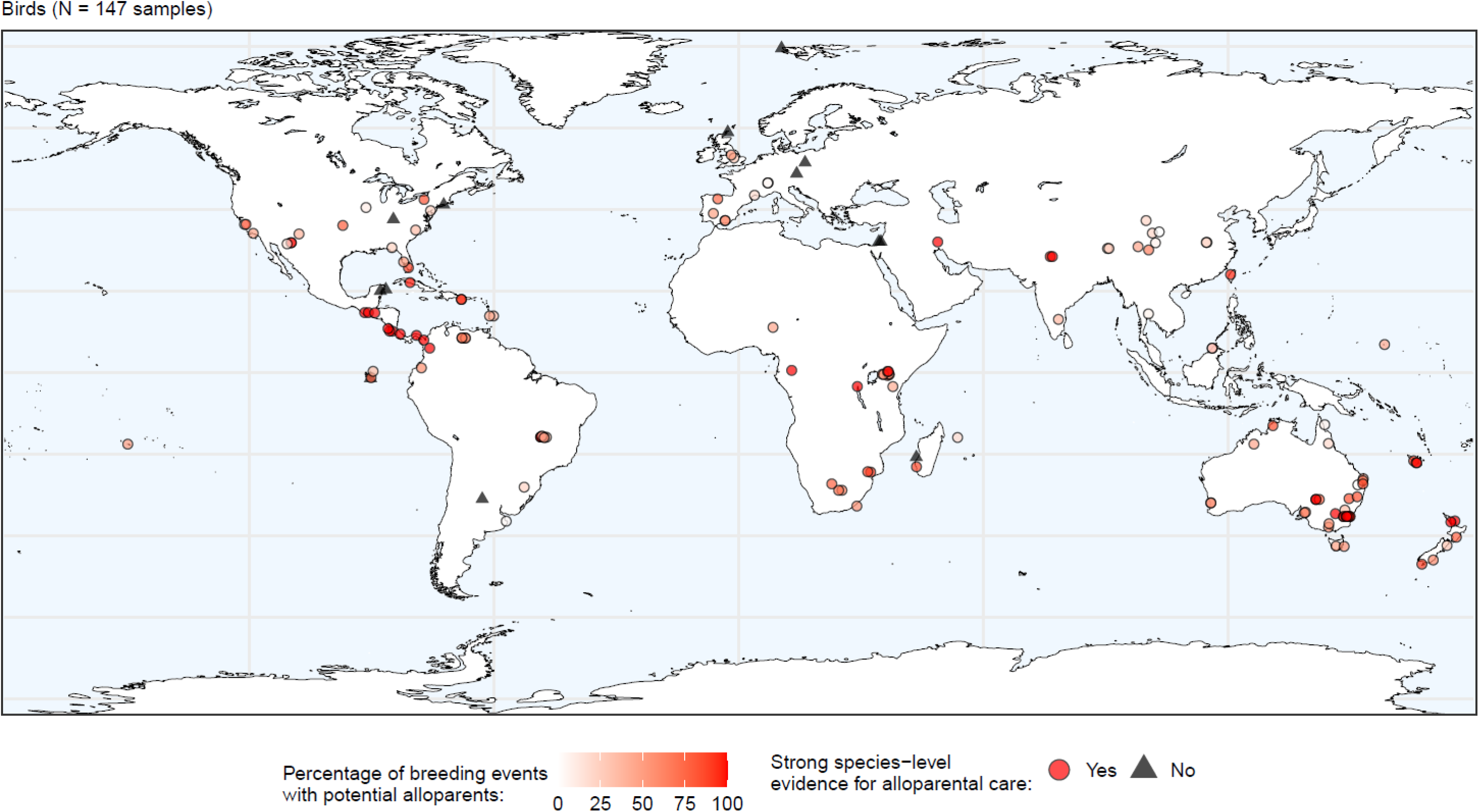
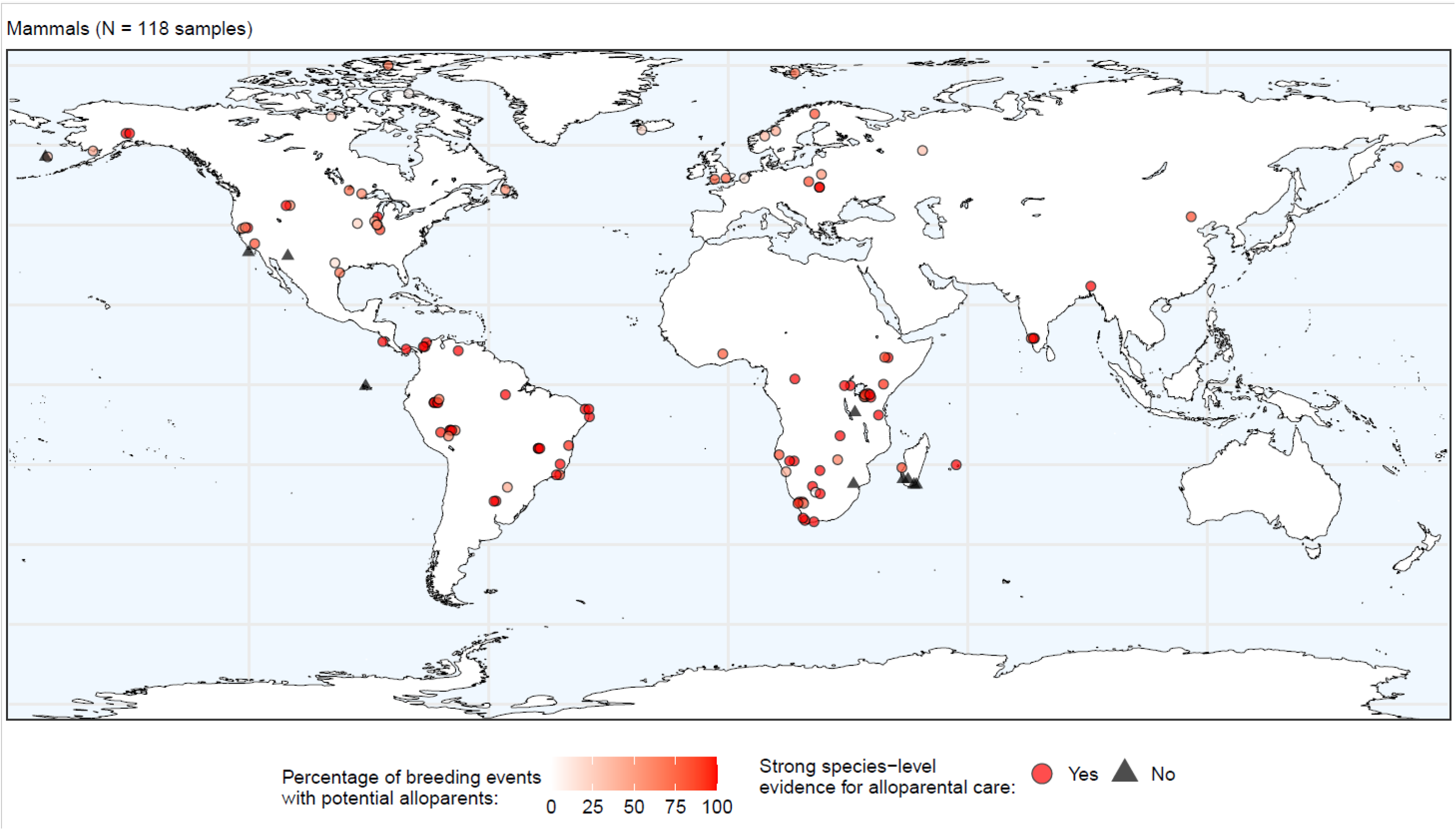
The geographical distribution of bird and mammal samples included in the Co-BreeD. *Prevalence of Alloparents* dataset. Each shape represents a sample from a specific sampling location and period.

#### Directing research effort

Estimates of breeding events with alloparents in mammals are often based on small sample sizes that hamper the drawing of solid conclusions (median: 12 breeding events, range: 1 – 3,234), especially when compared with birds (median: 54 breeding events, range: 1 – 2,792; Figure 2). Allocating resources to increase sample size is thus needed to ensure reliable inference, especially in cooperatively breeding mammals.

#### Intraspecific variation

The multi-year range of breeding events with alloparents is available for 126 samples from 84 species (Figure 1). Estimates of breeding events with alloparents from multiple populations are available for 45 species (range: 2-9 populations per species; Figure 1). In addition, the Co-BreeD *PA* appendix includes 70 year-by-year datasets from 47 species.

These detailed data enable (i) investigation of intraspecific variation across time and space, (ii) estimating the sampling error probability of samples (See ESM for demonstration of power analysis using Co-BreeD data.), and (iii) as well as testing the effect of selecting a single population per species for comparative analyses. Specifically, six (13%) of the 45 species with estimates from multiple populations include populations both under and above the 5% threshold of breeding units with alloparents (ground tit *Pseudopodoces humilis*, Australian magpie *Gymnorhina tibicen*, carrion crow *Corvus corone corone*, greater rhea *Rhea americana*, Arctic fox *Vulpes lagopus*, yellow mongoose *Cynictis penicillata*; see also Figure 1). These intraspecific variation data emphasise the importance of conducting comparative analyses with population-level rather than species-level data.

#### Cooperative breeding is more prevalent than previously recorded

The care systems of 88 bird and 55 mammal species classified in Co-BreeD were also included in other datasets on birds (Downing *et al*., 2020) and mammals (Lukas & Clutton-Brock, 2012) (Table 4). Notably, Co-BreeD reports evidence for alloparental care (see SMD comments for each species) for 62% of the bird (total N = 21) and 75% of the mammal species (total N = 20) classified by Downing *et al*. (2020) and Lukas & Clutton-Brock (2012) as non-cooperative or non-communal breeders, respectively. This considerable difference demonstrates the importance of reviewing data from multiple populations per species and of curating databases that can be corrected and updated with new natural history data.

**Table 4.** The number of species classified in Co-BreeD as having species-level evidence for cooperative breeding (i.e. species exhibits systematic alloparental care in >5% of breeding events in ≥1 population) and species classified as communal/cooperative breeders in a dataset on bird (Downing *et al*., 2020) and mammal care systems (Lukas & Clutton-Brock, 2012). Only species included in Co-BreeD and one of these two datasets are included. Disagreements are highlighted in red.

| Birds |  |  |  | Mammals |  |  |  |
| --- | --- | --- | --- | --- | --- | --- | --- |
|  |  | Co-BreeD |  |  |  | Co-BreeD |  |
|  |  | Cooperative breeder | Non-cooperative breeder |  |  | Cooperative breeder | Non-cooperative breeder |
| Downing *et al.* 2020 | Cooperative breeder | 66 | 1 | Lukas & Clutton-Brock 2012 | Cooperative breeder | 34 | 1 |
|  | Non-cooperative breeder | 13 | 8 |  | Non-cooperative breeder | 15 | 5 |

In total, Co-BreeD presents evidence for cooperative breeding (i.e. systematic alloparental care in >5% of breeding events) in at least one population of 31 species whose care system was classified as non-cooperative or were not included in recent datasets on the care systems of birds (Downing *et al*., 2020) and mammals (Lukas & Clutton-Brock, 2012). These include 16 bird (e.g. *Apteryx australis, Catharus fuscescens, Colius striatus, Cyanolyca argentigula, Cyanoramphus saisseti, Melanerpes erythrocephalus, Phoenicurus ochruros, Phyllastrephus placidus, Rhea americana, Rhynochetos jubatus, Sapayoa aenigma, Stachyris nigriceps, Staphida everetti, Sula sula, Vanellus chilensis*) and 15 mammal species (e.g. *Callithrix flaviceps, Cebuella pygmaea, Cebus capucinus, Cuon alpinus, Cynictis penicillata, Homo sapiens, Physeter macrocephalus, Saguinus fuscicollis, Saguinus geoffroyi, Sapajus nigritus, Tadarida brasiliensis, Trachypithecus pileatus, Vulpes macrotis, Vulpes vulpes*). Taken together, Co-BreeD results suggest that cooperative breeding is considerably more prevalent among birds and mammals than is currently appreciated.

### III.4. Adequate usage of the *PA* dataset

We make two suggestions to facilitate adequate use of the *PA* dataset. First, the *PA* parameter should not be used as a proxy for the proportion of breeding pairs versus groups in a population. This is because species that do not exhibit alloparental care are indicated as having 0% of breeding events with alloparents, although they often live in groups during the breeding season. For instance, white-breasted mesites (*Mesitornis variegatus*) and Siberian jays (*Perisoreus infaustus*) often breed in family groups, but only parents care for their offspring (Gamero, Székely & Kappeler, 2014; Ekman & Griesser, 2016).

Second, the *PA* parameter is independent of the extent of alloparental care provided to offspring. Hence, species in which most offspring receive alloparental care may have a high percentage in the *PA* parameter even if the extent of alloparental care is limited. These cases are especially problematic in group-living species. For example, ring-tailed lemurs almost always live in groups where offspring receive very little alloparental care (Gould, 1992; Sauther, Sussman & Gould, 1999) and the species is nevertheless classified as 100% of breeding units having potential alloparents. Users of the *PA* dataset are thus advised to carefully use those estimates where the prevalence of breeding events with potential alloparents was assessed based on group size greater than two adults rather than on observed alloparental care (see Table 2). This problem will be minimised with the expansion of Co-BreeD to include parameters on different alloparental care behaviours, and thereby, establishing a more comprehensive identification index for cooperatively breeding species.

## IV. Conclusions

1. The Cooperative-Breeding Database (Co-BreeD) is an open-access, peer-reviewed and updateable resource that covers key biological parameters of cooperatively breeding species.
2. It has a sample-based structure where each sample is linked to a sampling location and period, enabling the (i) retrieval of time and space-specific environmental data, and (ii) exploration of within-species variability in cooperative breeding behaviours (Baglione *et al*., 2002; Lu *et al*., 2011 and Figure 1).
3. Co-BreeD appendices archive datasets with year-by-year resolution covering a total of 384 research years on 47 species. This documentation includes many unpublished datasets. The archiving of these unpublished datasets is particularly important for the preservation of natural history behavioural data as habitat loss also results in loss of behavioural diversity (Kühl *et al*., 2019).
4. By providing extended data that include percentage estimates, sample sizes and year-by-year breakdowns, Co-BreeD enables the estimation of additional statistical parameters; for example, the probability of sampling error associated with each value (see ESM) and weighted analyses according to sample sizes.
5. The continuous measure of the prevalence of breeding events with alloparents advances cooperative breeding research beyond the traditional binary species classification (i.e. cooperative vs. non-cooperative breeders) towards analysing cooperative breeding as a continuous trait. This approach significantly increases the explanatory power of statistical analyses (Clutton-Brock, 2021; Olivier *et al*., 2024).
6. The data from the Co-BreeD Prevalence of Alloparents dataset suggests that cooperative breeding in birds and mammals is more prevalent than currently estimated.
7. Co-BreeD focuses on species with actual data about the care system. However, since reliable data on alloparental care is currently missing for most species (Cockburn, 2006; Lukas & Clutton-Brock, 2017; Schradin, 2017), Co-BreeD is set as a living resource that can be expanded with the publication of new data. It is therefore an ongoing collaborative effort of many members of the cooperative breeding research community and we encourage researchers to propose corrections, contribute from their data and join the curating effort of Co-BreeD.

## Authors contribution

Research design: Y.B.M., S.M.D., M.G.

Funding acquisition: Y.B.M., S.M., M.G.

Data collection: Y.B.M., M.W.

Contribution of unpublished datasets to Co-BreeD: V.B., L.C., R.C., C.D., R.G., J.L., K.M.M., J.T., D.A.W.

Data analysis: Y.B.M., S.M.D.

Data validation: Y.B.M., M.W., S.M., F.F., V.B., J.B., L.C., R.C., G. H. B. d. M., C.J.D., C.D., R.G., R.H., S.A.K., J.L., K.M.M., A.N.R., C.R., D.R.R, C.S., J.T., M.H.W., D.A.W., I.A.W.

Visualisation: Y.B.M., F.F.

Writing of manuscript: Y.B.M.

Editing of the manuscript: Y.B.M., M.W., S.M.D., S.M., F.F., V.B., J.B., L.C., R.C., G. H. B. d. M., C.J.D., C.D., R.G., R.H., S.A.K., J.L., K.M.M., A.N.R., C.R., D.R.R, C.S., J.T., M.H.W., D.A.W., I.A.W., M.G.

